# Full Atomistic Model of Prion Structure and Conversion

**DOI:** 10.1101/505271

**Authors:** Giovanni Spagnolli, Marta Rigoli, Simone Orioli, Alejandro M. Sevillano, Pietro Faccioli, Holger Wille, Emiliano Biasini, Jesùs R. Requena

## Abstract

Prions are unusual protein assemblies that propagate their conformationally-encoded information in absence of nucleic acids. The first prion identified, the scrapie isoform (PrP^Sc^) of the cellular prion protein (PrP^C^), is the only one known to cause epidemic and epizootic episodes(1). Most aggregates of other misfolding-prone proteins are amyloids, often arranged in a Parallel-In-Register-β-Sheet (PIRIBS)(2) or β-solenoid conformations(3). Similar folding models have also been proposed for PrP^Sc^, although none of these have been confirmed experimentally. Recent cryo-electron microscopy (cryo-EM) and X-ray fiber-diffraction studies provided evidence that PrP^Sc^ is structured as a 4-rung β-solenoid (4RβS)(4, 5). Here, we combined different experimental data and computational techniques to build the first physically-plausible, atomic resolution model of mouse PrP^Sc^, based on the 4RβS architecture. The stability of this new PrP^Sc^ model, as assessed by Molecular Dynamics (MD) simulations, was found to be comparable to that of the prion forming domain of Het-s, a naturally-occurring β-solenoid. Importantly, the 4RβS arrangement allowed the first simulation of the sequence of events underlying PrP^C^ conversion into PrP^Sc^. Our results provide the most updated, experimentally-driven and physically-coherent model of PrP^Sc^, together with an unprecedented reconstruction of the mechanism underlying the self-catalytic propagation of prions.

**Significance:** Since the original hypothesis by Stanley Prusiner, prions have represented enigmatic agents diverging from the classical concept of genetic inheritance. However, the structure of PrP^Sc^, the infectious isoform of the cellular prion protein (PrP^C^), has so far remained elusive, mostly due to technical challenges posed by its aggregation propensity. Here, we present a new high resolution model of PrP^Sc^ derived from the integration of a wide array of recent experimental constraints. By coupling the information of such model with a newly developed computational method, we reconstructed for the first time the conformational transition of PrP^C^ to PrP^Sc^. This study offers a unique workbench for designing therapeutics against prion diseases, and a physically-plausible mechanism explaining how protein conformation could self-propagate.

## Introduction

Prion diseases are infectious neurodegenerative disorders characterized by an invariably lethal outcome caused by a proteinaceous infectious agent named “prion”(1). The central event in these pathologies is the conversion of PrP^C^, a GPI-anchored protein of unknown function, into a misfolded isoform (PrP^Sc^) which accumulates in the central nervous system of affected individuals(6). While PrP^C^ structure has been widely characterized, and consists of a N-terminal disordered tail and a C-terminal globular domain(7), no high-resolution information is available for PrP^Sc^ due to technical challenges posed by its high insolubility and aggregation propensity(8). In order to fill this gap, different atomistic models based on low-resolution experimental data have been proposed, including a Left-handed-β-Helix (LβH) structure spanning residues 89 to 170 while retaining the two C-terminal α-helices of PrP^C^ (9), and a Parallel In-Register Beta-Sheet (PIRIBS) architecture, characterized by intermolecular stacking of aligned PrP monomers(10). The PIRIBS model is not consistent with recent cryo-EM data obtained using infectious, anchorless PrP^Sc^ fibrils (Supp. Fig. 1)(4). Moreover, this model fails to accommodate glycosylated residues in PrP^Sc^, which would result in the introduction of excessive steric clashes(11). In turn, while consistent with the mentioned experimental constraints, the proposed LβH model is incoherent with a recent re-evaluation of previous FTIR data suggesting that PrP^Sc^ does not contain α-helices(8).

**Fig. 1.**
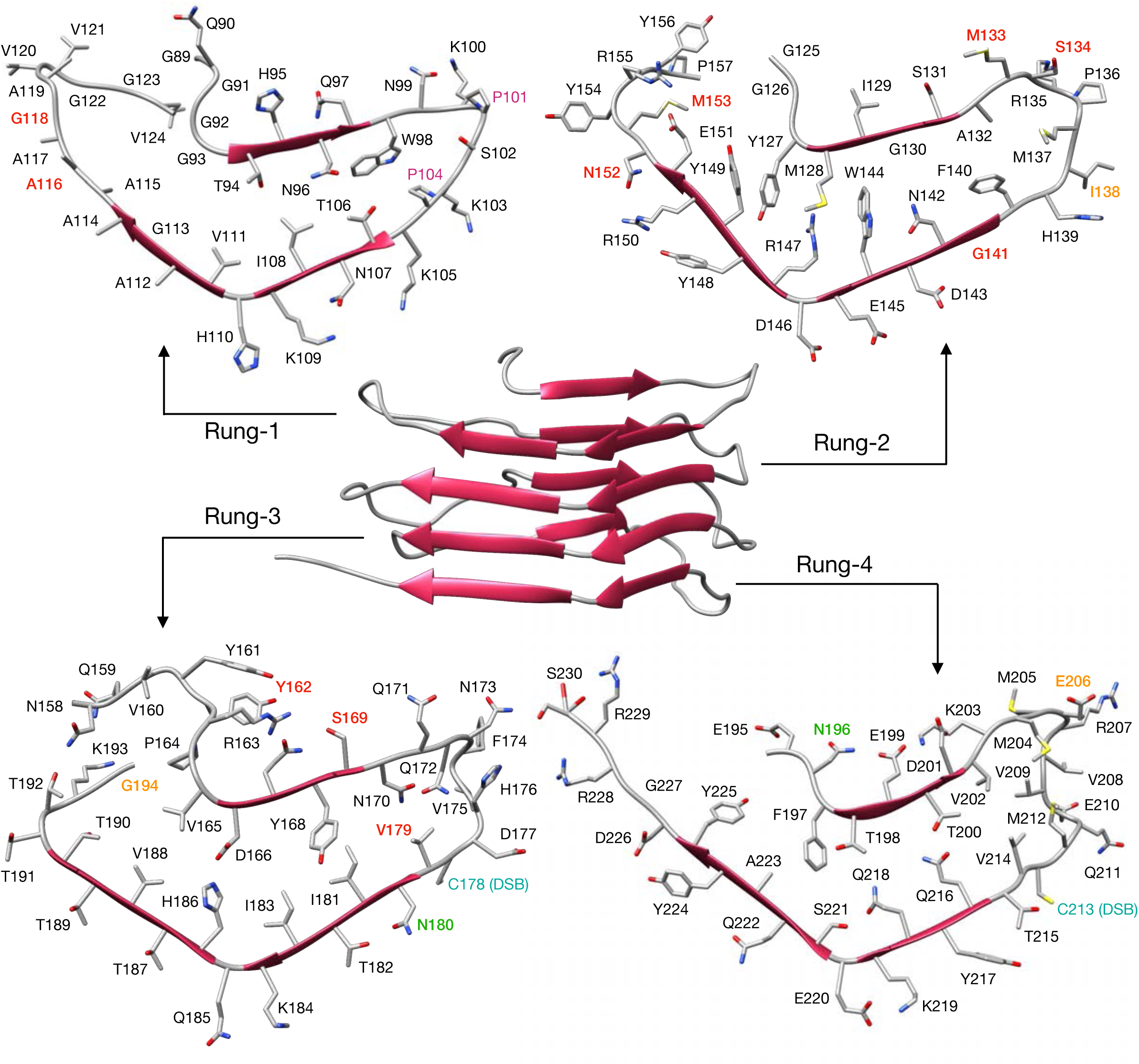
View of the 4RβS PrP^Sc^ model. The structure of PrP^Sc^ modelled as a 4RβS (β-strands represented as red arrows) is depicted in the center of the figure. Residues are displayed in each individual rung (1–4) with different colors. PK cleavage sites identified by mass spectrometry in two different previous reports are colored in red (13) or orange(14). Glycosylation sites are labelled in green. Prolines are colored in purple. Cystine is indicated in cyan.

## Results

To satisfy current experimental evidence and theoretical structural constraints, we built a new atomistic model of mouse PrP^Sc^ based on the 4RβS conformation, and tested its stability by means of all-atom MD simulations. The construction of the model took into account an array of experimental data, including: (i) cryo-EM(4) and X-ray fiber-diffraction studies(5), which showed that the fold of a mouse glycosylphosphatidylinositol (GPI)-anchorless, infectious PrP^Sc^ is compatible with a 4RβS architecture with L-or T-shaped cross-section(4); (ii) circular dichroism (CD) and FTIR spectroscopy, which ruled out the presence of α-helices, and suggest that PrP^Sc^ contains approximately 40-50% β-sheet and 50-60% coil/turns(8); (iii) Mass Spectrometry (MS) analyses indicating the presence of an intact disulphide bond between residues C178 and C213 (mouse sequence)(12), as well as mapping Proteinase K (PK)-sensitive residues, which reflect amino acids likely excluded from the resistant core of the protein(13),(14); and (iv) the possibility of accommodating complex glycans at positions N180 and N196(11). All these constraints were comprehensively included into a 2D threading scheme spanning mouse PrP (moPrP) residues 89-230 (Supp. Fig. 2), also considering the structural propensities of different residues: polyglycine tracts and prolines were positioned in loops due to their destabilizing effects on β-strands; charged sidechains were excluded from the inner core of the protein or counterbalanced by salt bridges. This scheme was then modelled onto the 3D arrangement of a naturally-occurring β-solenoid protein (*Dickeya dadantii* Pectate Lyase; PDB 1AIR). The resulting structure (depicted in Fig. 1) features an inner core containing mainly hydrophobic or mildly polar side-chains (T94, T106, L108, V111, Y127, M128, W144, Y149, V165, Y168, I181, I183, V188, F197, T198 and T200), few polar side-chains involved in hydrogen bonding (N142-HB-Y168, H168-HB-T198, Q216-HB-T200 and Q218-HB-S221), and a salt bridge (R147-SB-D166). Conversely, the majority of the highly-polar residues (N and Q) including the glycosylation sites (N180 and N196) and charged side-chains (E, D, K and R) are exposed to the solvent. The structure also encompasses identified PK cleavage sites localized in loops/turns, or at the edge of the β-strands, and the intact disulphide bond between C178 and C213. Importantly, the final model fitted with a previously described, low-resolution cryo-EM map of infectious PrP^Sc^ (Supp. Fig. 3).

**Fig. 2.**
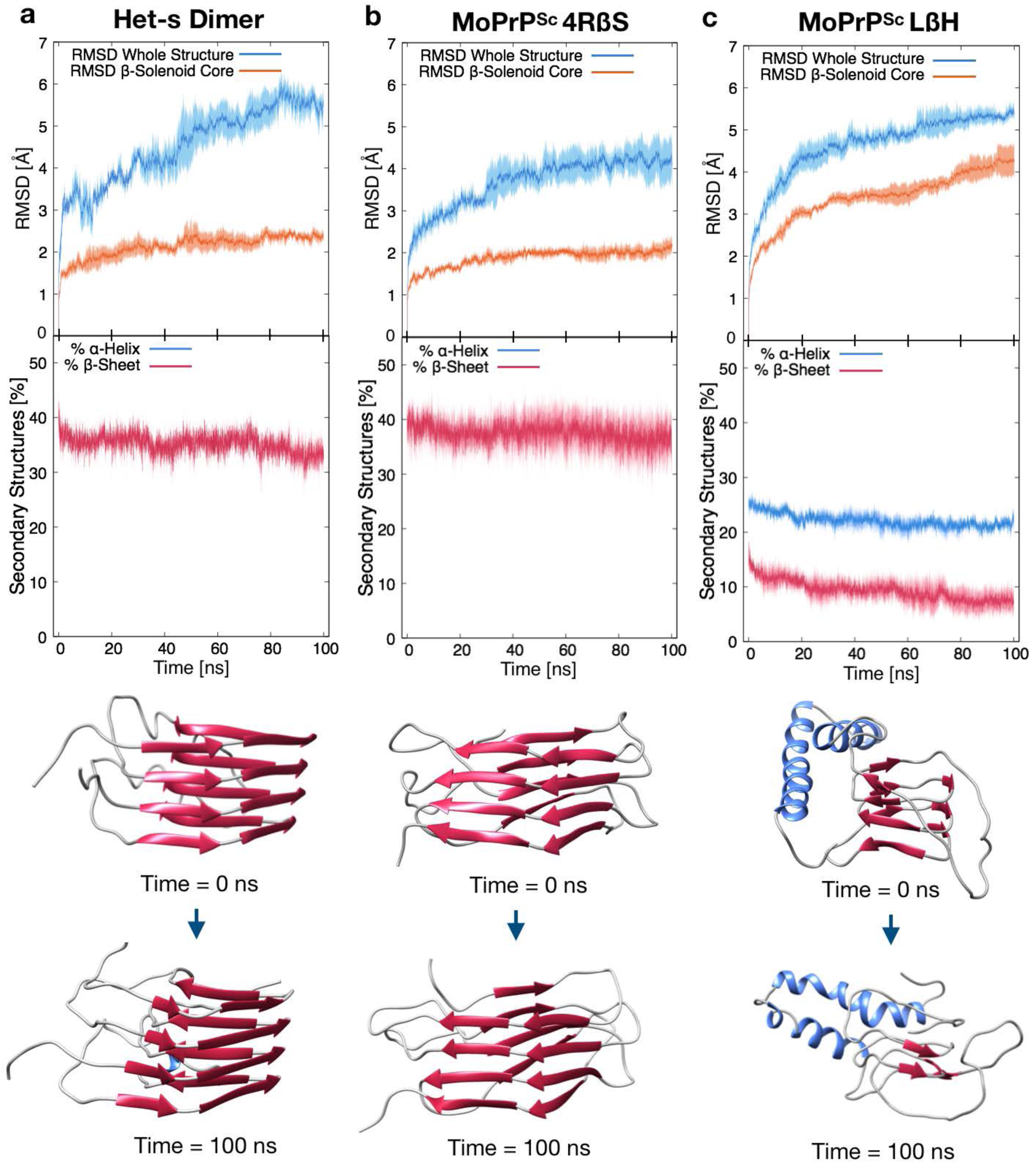
Comparison of stability by MD simulations of a Het-s dimer, 4RβS and LβH PrP^Sc^. Upper graphs report the RMSD deviation from initial configuration for the entire structure (blue lines) or the β-solenoid core (orange lines) of the different proteins. Filled curves indicate standard error of the mean. Results show a comparable stability between the Het-s dimer (a) and the 4RβS model (b), with an average RMSD of the hydrophobic core (calculated as the average of the three trajectories over the last 5 ns, ± standard deviation) of 2.4 ± 0.2 Å for the Het-s dimer (similar results for Het-s stability are reported(30)) and 2.1 ± 0.3 Å for the 4RβS. In contrast, the structural deviation of the LβH (c) hydrophobic core is approximately two-fold higher, reaching a value of 4.3 ± 0.6 Å. Lower graphs indicate the α-helical (blue lines) or β-sheet (red lines) content of each protein. The initial and final β-sheet content was calculated as the average of the three trajectories over the last 5 ns of the restrained and unrestrained MD simulations, respectively. The Het-s dimer showed a variation from an initial 41.8 ± 2.3% to a final 33.2 ± 2.8%. Similarly, the 4RβS model deviates from an initial 41.3 ± 2.3% to a final 36.6 ± 5.4%. Instead, the LβH model deviates from a starting 18.3 ± 1.4% to a 7.5 ± 3.1%. These results are illustrated by the structures shown below the graphs, which represent the initial (top) and final (bottom) frames of the MD trajectories.

**Fig. 3.**
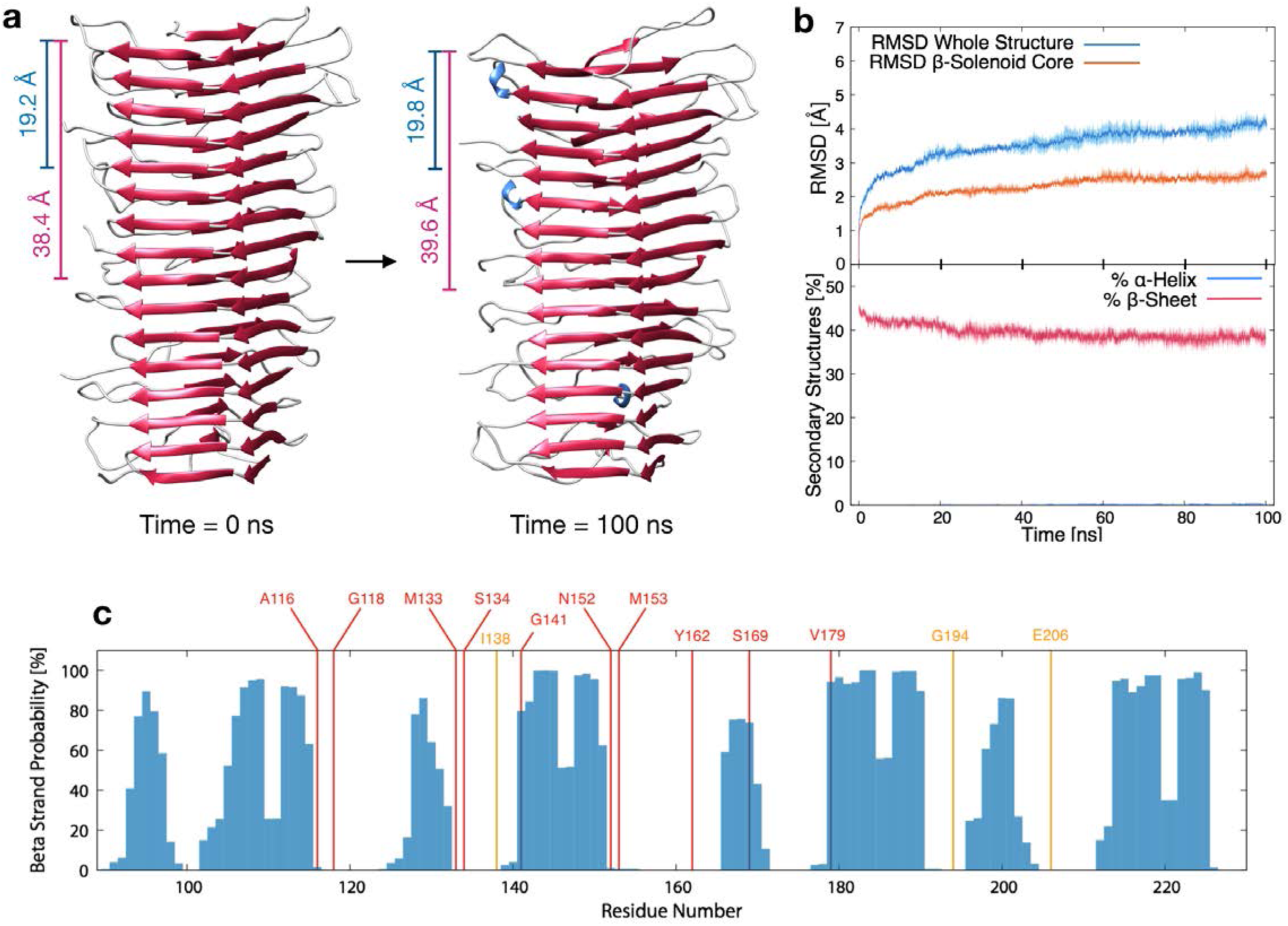
MD simulation and PK restriction map of a 4RβS tetramer. **a,** 3D representation of a 4RβS tetramer at the beginning (left) and the end (right) of MD simulations. Blue bars indicate the distance between two residues in the same position on two consecutive monomers, which corresponds to 19.2 ± 0.4 Å (t = 0 ns) and 19.8 ± 2.2 Å (t = 100 ns). Purple bars indicate the distance between a residue in one monomer and the same residue on the second forthcoming monomer, which corresponds to 38.4 ± 0.5 Å (t = 0 ns) and 39.6 ± 1.8 Å (t = 100 ns). A similar pattern of signals reflecting monomeric and dimeric repeats has previously been observed by cryo-EM studies on Het-s(31). Both values are in almost perfect agreement with the two main signals obtained by Fourier transform single particle analysis in the cryo-EM experiment (19.1 Å and the ~40 Å signals). The average monomeric model volume calculated at beginning and at the end of the 100 ns tetramer simulations are equal to (18.5 ± 0.7)•10^3^ Å^3^ and (18.5 ± 0.3)Å10^3^ Å^3^ respectively; while the estimated cryo-EM monomeric volume from the fibril is equal to 18.9Å10^3^ Å^3^ **b**, Upper graph shows the RMSD deviation of the tetramer from initial state for the entire structure (blue lines) or the β-solenoid core (orange lines). Structural deviation over the 100 ns of simulation corresponds to 2.6 ± 0.2 Å. Lower graph report the secondary structures percentage, initial β-strand content is 46.2 ± 1.2 %, while the final 38.6 ± 1.7%. Filled curves indicate standard error of the mean. These results indicate a comparable stability between monomeric and tetrameric 4RβS structures. **c**, Map of the experimentally observed PK cleavage sites (identified by mass spectrometry in two previous reports are colored in red or orange) overlapped with the probability of each residue to be in an extended conformation, calculated over the last 5 ns of the MD trajectories.

To test the physical consistency of the 4RβS model, we challenged its stability by all-atom MD simulations in explicit solvent. First, three independent, 20 ns simulations were performed in the isothermal-isobaric ensemble (T = 300 K, P = 1 Bar) restraining hydrogen bonds distances between atoms involved in β-sheets. This process allowed relaxation of protein loops and side chains of the core (Supp. Fig. 4). Next, the imposed restraints were released, and three plain-MD trajectories of 100 ns each were simulated. The stability of the model was then assessed by both Root Mean Squared Deviation (RMSD) of atomic positions relative to the initial frame (t = 0 ns) and secondary structures content. Interestingly, we obtained values in the same range of fluctuation for our 4RβS model and the prion-forming domain of the fungal protein Het-s, a naturally occurring β-solenoid whose structure has been solved by solid-state NMR (ssNMR; PDBs 2KJ3 and 2RNM; Fig. 2, and Supp. Fig. 5). Conversely, by applying an identical workflow to the previously proposed model of laterally-stacked LβH trimer of PrP^Sc^, we observed a profound instability of the β-helical domain, which was already evident after few tens of ns of dynamics (Fig. 2 and Supp. Fig. 5 and 6).

**Fig. 4.**
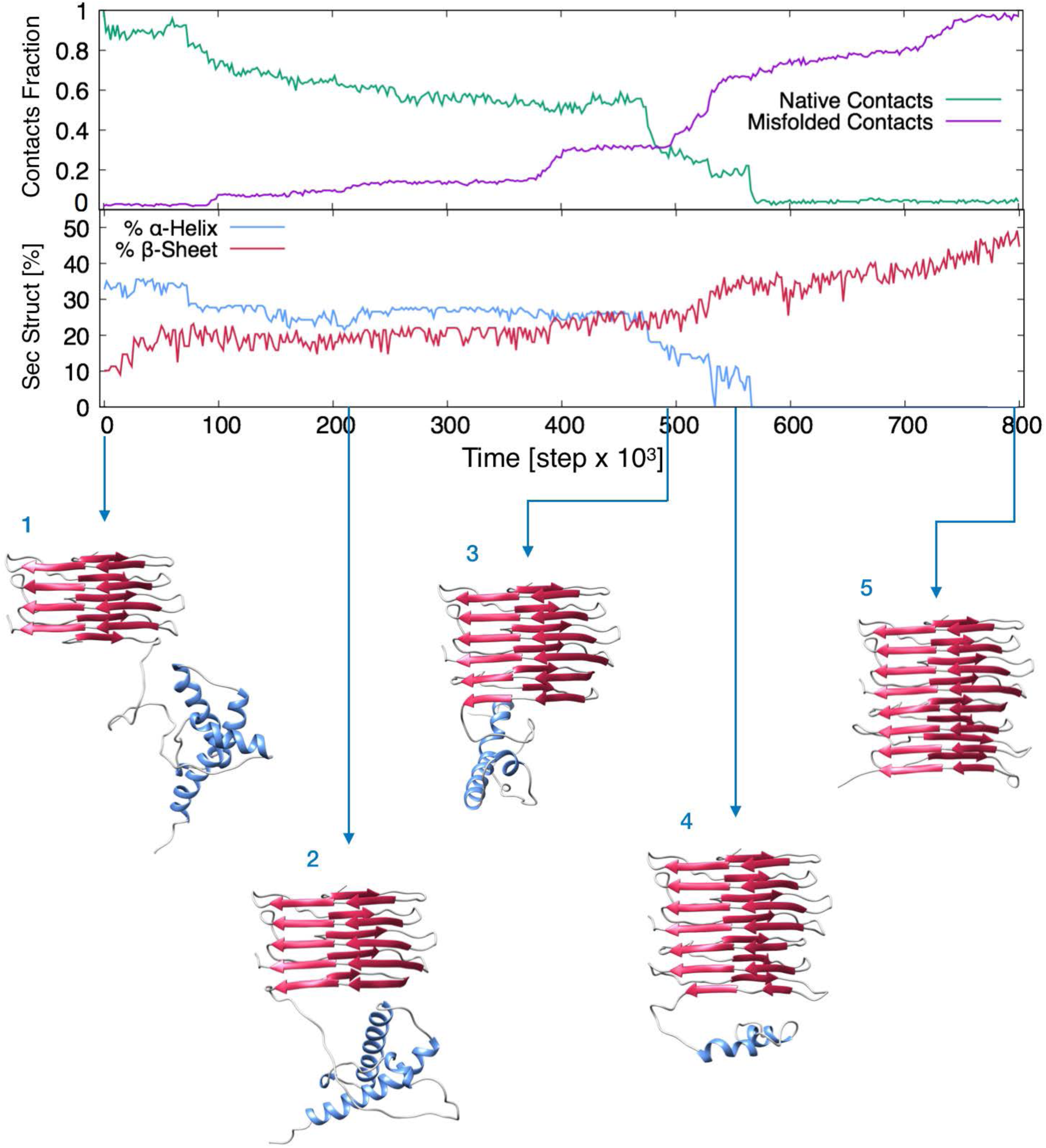
Graphs and frames extracted from the rMD simulation of PrP^C^ to PrP^Sc^ conversion. Upper graph reports the fraction of native and misfolded contacts of the instantaneous configurations of the PrP^C^-PrP^Sc^ complex, starting from the initial modelled configuration (depicted in 1) in which PrP^C^ contacts the 4RβS monomer, with respect to the final target configuration (4RβS dimer). Lower graph shows the secondary structure content. Pictures of the evolving complex represent frames extracted from the entire conversion simulation at precise rMD steps, corresponding to the refolding of PrP^C^ as follow: (2) residues 89-115; (3) residues 89-151; (4) residues 89-190; (5) residues 89-230. The process highlights the progressive unfolding and refolding of PrP^C^ onto the 4RβS template, which initially involves the unstructured region, followed by the loss of α-helices and a progressive formation of β-sheets.

To mimic the fibrillary behavior of PrP^Sc^, we built a tetrameric 4RβS model by stacking monomers in a head-to-tail fashion. As expected, this assembly showed a comparable MD stability to the monomer (Fig. 3). Moreover, the 4RβS tetramer fits with the two main signals obtained by Fourier transform single particle analysis of cryo-EM data (19.1 Å and the ~40 Å)(4), which reflect distances between the same residues on two contiguous or alternate monomers, respectively, as well as with the observed volume of the fibrils (Fig. 3). Finally, by introducing complex sugar precursors (GlcNAc2Man3Fuc) at positions N180 and N196 of each monomer, we observed the absence of steric clashes, confirming that the 4RβS model accommodates the presence of glycans (Supp. Fig. 7). Collectively, these findings indicate that, in contrast to the LβH model, the 4RβS is a solid arrangement for PrP^Sc^, built on the most recent experimental data, and showing a conformational stability comparable to that of a natural prion.

The 4RβS model allowed us to develop an original scheme to perform for the first time a simulation of the conformational transition from PrP^C^ to PrP^Sc^. In order to bridge the gap between the computationally-accessible and the biologically-relevant time scales, we employed a specific kind of biased dynamics called ratchet-and-pawl MD (rMD)(15) in the framework of an all-atom realistic force field(16). rMD-based methods have been successfully applied to simulate protein folding and other conformational transitions(17, 18). However, this scheme provides a sampling of the transition path ensemble only if the biasing force is applied along a reliable reaction coordinate (19). Therefore, the first step towards developing our rMD simulation was to build a statistical model to identify the reaction coordinate of the process. Using the Markov State Mode formalism, we demonstrate that among all the possible misfolding patterns from PrP^C^ to PrP^Sc^ the prominent reaction mechanism is the sequential formation of rungs. A biasing coordinate was then built by coupling this dynamical information with the all atom 4RβS structure. The associated rMD simulations yielded a transition pathway with full atomistic resolution in which the C-terminal rung of the solenoid acts as a primary conversion surface for PrP^C^ unstructured N-terminus (residues 89-124). This first event initiates a cascade of conformational transitions in which each newly formed rung acts as a template for the formation of the following one, ultimately leading to the complete conversion of PrP^C^ into PrP^Sc^ (Fig. 4 and Supp. Movie). This analysis provides a rigorous description of the active role of fibril ends in the templated propagation of prions, and it is compatible with previous observations suggesting the presence of a structured intermediate conformer of PrP^C^ in its transition to PrP^Sc^ (20)

## Discussion

The search for an effective therapy against prion diseases has largely been unsuccessful, partly due to the lack of information regarding the structure of PrP^Sc^, which has hampered rational drug discovery efforts. Indeed, the elucidation of the structure of PrP^Sc^ at atomic resolution has proven to be a phenomenal experimental challenge, mainly due to its high insolubility and aggregation propensity. Available computational models of PrP^Sc^ have the virtue of providing a plausible 3D structure, but fail to comprehensively accommodate most recent experimental data. Here, we filled this gap by exploiting the information arising from cryo-EM and X-ray fiber-diffraction studies, which suggested a general architecture as a 4RβS, and refined the structure by including experimental constraints obtained by mass spectrometry. This new model possesses several important features. It displays an intrinsic coherence with *state-of-the-art* knowledge about infectious PrP^Sc^ fibrils, and it appears to be as stable as a naturally-occurring prion. It also represents a unique workbench for interpreting future structural data or available biological evidence, such as the effect of variations in the *PRNP* gene favoring or disfavoring prion propagation. Importantly, the 4RβS model allowed us to perform the first reconstruction of how the information encoded into the conformation of a protein could be propagated in a directional fashion, a concept underlying the infectious nature of prions.

## Methods

### Template Selection & Model Building

In order to accommodate the moPrP 89-230 sequence, we selected a β-solenoid architecture capable of satisfying the following requirements: (i) Approximate number of residues in extended conformation higher than 12 per rung. Requirement needed to fit the secondary structure content of approximately 40% of β-sheets(8); (ii) Capacity of accommodating two extended loops per rung. Namely, the possibility of introducing arbitrarily long sequences between consecutive β-strands in order to allow the exclusion of PK cleavage sites, glycine tracts and prolines from the core of the solenoid; (iii) L or T cross-section solenoidal-shape. Architecture coherent with cryo-EM maps; (iv) Capacity of accommodating bulky aromatic residues in the hydrophobic core (e.g. moPrP 89-230 contains 11 Tyr, 3 Phe and 1 Trp). All these requirements are satisfied by the right-handed, L-shaped β-solenoid architecture. In contrast, left-handed-solenoid structures typically display smaller rungs(21), impairing the modelling of the desired number of residues in extended conformation as well as the accommodation of bulky residues in the core. The left-handed structure of the prion forming domain of Het-s (PDB 2KJ3 and 2RNM) is an exception in terms of β-sheet content, however its hydrophobic core is mainly composed of small side-chains and it allows the insertion of only one arbitrarily long loop connecting two consecutive rungs(3). The threading scheme for the PrP 89-230 sequence (Supp. Fig. 2) was obtained by following the L-shaped β-solenoid architecture. In the threading process, PK cleavage sites, prolines and glycine tracts were positioned in the loops (when this was not possible, at the edges of beta strands). Charged side-chains were excluded from the inner core of the solenoid or paired with residues forming salt bridges. The presence of an intact disulphide bond between C178 and C213 and solvent exposure of N180 and N196 sidechains (in order to accommodate glycans) were also considered.

The 3D structure of the monomeric model of PrP^Sc^ was constructed following these steps: (i) 4 Rungs of *Dickeya dadantii* Pectate Lyase (PDB 1AIR; two repetitions of residues 168-235) were used to obtain the scaffold for the β-solenoid core; (ii) The original loops of the protein were removed; (iii) The residues in the hydrophobic core of the original PDB structure were replaced with PrP residues (as indicated in Supp. Fig. 2) using Chimera(22); (iv) New loops were generated using the MODELLER tool(23) in Chimera. Each loop was selected from a set of 20 proposed conformations. Structures resulting in extended atomic clashes were discarded, and finally the best performing model in term of DOPE score was selected; (v) Side-chain geometry was optimized using the Dunbrack’s rotamer library. In particular, for highly polar and charged side-chains in the hydrophobic core, geometries capable of forming hydrogen bonds or salt bridges with nearby residues were selected; (vi) The energy of the system was minimized in vacuum with the steepest descent algorithm in Gromacs 4.6.5(24). System topology was generated using Amber99SB-ILDN force field(16). A restraining potential during energy minimization between the H and O atoms involved in hydrogen bond formation between backbone residues was added, as in equation 1:

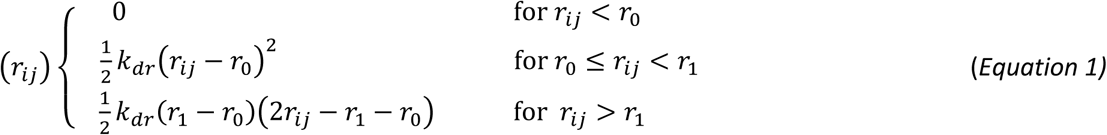

Where r_ij_ is the distance between the H and the O atoms involved in hydrogen bond formation; *r*_0_ is the original hydrogen bond distance after energy minimization (2 < r_0_ < 2.5) Å and *r*_1_ is an upper limit equal to 4 Å, while *k*_dr_ is set to 2⋅10^3^ kJ/(mol⋅nm^2^).

This strategy was applied to favor the movement of the side-chains and accommodation of loops, while impairing the backbone deviation of the residues involved in the β-solenoid core formation; (vi) Additional optimization of the backbone (in order to remove Ramachandran outliers) and side-chain geometry was performed using Coot(25); (vii) Absence of steric clashes was verified using Chimera, setting the VDW-overlap threshold as 0.6 Å, subtracting 0.4 Å to account H-Bonding pairs and ignoring contacts of pairs <4 bonds apart. The 3D structure of the tetrameric model of PrP^Sc^ was assembled by stacking four monomers in a head-to-tail fashion using Chimera, maintaining the proposed threading. The strands 198-TETD and 215-TQYQKESQAYY were stacked on the top of 94-THNQ and 105-KTNLKHVAGAA of the forthcoming monomer, respectively. The structure was then energy minimized following the same protocol used for the monomer. Glycans (DManpα 1-6[DManpα 1-3]DManpβ 1-4DGlcpNAcβ 1-4[LFucpα 1-6]DGlcpNAcα 1-Asn) attached to residues N180 and N196 were added using the doGlycans tool to the 4RβS tetramer, and the structure was energy minimized in vacuum and then explicit solvent using Gromacs 5.1.2.

### Molecular Dynamics Simulations

Molecular Dynamics simulations were performed on the supercomputers Finis Terrae II (CESGA, Santiago de Compostela, Spain) and Marconi (CINECA, Bologna, Italy) using Gromacs 5.1.2. The following protocol was applied for the whole set of simulations performed in this work. Protein topologies were generated using Amber 99SB-ILDN force field. The structures were accommodated in a dodecahedral box with periodic boundary conditions. The minimum wall distance from the protein was set to 12 Å. The box was filled with TIP3P water molecules. The charge of the system was neutralized with the addition of Na^+^ or Cl^-^ ions. The system was energy minimized in explicit solvent using a steepest descent algorithm, retaining the restraining potential (equation 1). From the minimized structure, three independent equilibrations with position restraints on heavy atoms were launched: in the NVT ensemble (300 K) for 500 ps using the Nose-Hoover thermostat, and then in the NPT ensemble (300 K, 1 Bar) for 500 ps using the Nose-Hoover thermostat and the Parrinello-Rahman barostat. For each equilibrated system, a 20 ns MD simulation with restraining potential (Equation 1) was launched. Finally, restraints were released and each trajectory was extended with 100 ns of plain-MD. This protocol yielded three independent, unbiased 100 ns MD trajectories for each structure of interest. MD simulations were performed using a leap-frog integrator with step equal to 2 fs. LINCS algorithm was applied to handle bonds constraints. Cut-off for short-range Coulomb and Van der Waals interactions was set to 12 Å, while long range electrostatics was treated using Particle Mesh Ewald.

### Generation of the Propagation Model

A key step to perform reliable rMD simulations is the identification of a reasonable reaction coordinate. To this end, we first developed a statistical coarse-grained model which enables us to identify the dominant reaction pathway underlying the conversion of PrP^C^ into a 4RβS PrP^Sc^. In order to describe the reaction kinetics we developed a simple stochastic model in which the key rate-limiting processes are assumed to be the irreversible formation and docking of the four rungs of the 4RβS. We indicated R_0_ as the C-terminal rung of the pre-formed 4RβS, and with R_1_ R_2_ R_3_ R_4_ the consecutive rungs of the converting monomer. The instantaneous state of the system can be represented as a 4-dimensional vector of binary entries S = [n_0_, n_1_, n_2_, n_3_], where n_k_ = 1 in the presence of docking between rung R_k_ and rung R_k+1_, and n_k_ = 0 otherwise. We emphasize that this model excludes the presence standalone rungs, which would correspond to an entropically-unfavorable single extended conformation, not stabilized by hydrogen bonds of nearby β-strands. On the other hand, misfolded rungs can stabilized either upon docking to a pre-existing misfolded region (template mechanism) or through a process in which two rungs simultaneously form and dock. We modeled the transition of an initial state S_R_ = [0,0,0,0] (in which the PrP^C^ monomer is in the native state and none of the rungs are formed) to the fully misfolded state S_P_ = [1,1,1,1] (where the monomer is completely misfolded and incorporated into PrP^Sc^). The resulting network is represented in Supp. Fig. 8a which contains all the possible combinations of docking events leading S_R_ to S_P_ through a sequence of irreversible transitions. The model can be simplified considering that rate k_2_ is expected to be negligible as compared to k_0_. Indeed, while the event associated with k_0_ only requires the structuring of a disordered PrP region, the events associated with k_2_ require the loss of native content (breakage of hydrogen bonds in the helical regions) together with the simultaneous formation of two rungs (two-fold entropic cost). Thus, it is possible to disregard all the reaction pathways in which the first step of the reaction is a transition occurring at a rate k_2_ and consider only events starting from [1,0,0,0]. The network of the resulting simplified model is depicted in Supp. Fig. 8b. Furthermore, we can set k_3_/k_1_ >>1 and k_3_/k_2_ >>1, since the docking of two pre-formed rungs occurs at a rate much faster than all processes involving misfolding. The reaction kinetics in this stochastic model was simulated through a Kinetic Monte Carlo algorithm (arbitrarily setting k_3_/k_1_ = 10^6^) and the resulting reaction mechanisms were studied as a function of the k_2_/k_1_ ratio. Namely, we enumerated all the reaction pathways and computed the corresponding probability:

|  | Step 1 | Step 2 | Step 3 | Step 4 |
| --- | --- | --- | --- | --- |
| Path 1 | [1,0,0,0] | [1,1,0,0] | [1,1,1,0] | [1,1,1,1] |
| Path 2 | [1,0,0,0] | [1,0,1,0] | [1,1,1,0] | [1,1,1,1] |
| Path 3 | [1,0,0,0] | [1,0,0,1] | [1,0,1,1] | [1,1,1,1] |
| Path 4 | [1,0,0,0] | [1,0,0,1] | [1,1,0,1] | [1,1,1,1] |
| Path 5 | [1,0,0,0] | [1,1,0,0] | [1,1,0,1] | [1,1,1,1] |
| Path 6 | [1,0,0,0] | [1,0,1,0] | [1,0,1,1] | [1,1,1,1] |

We find that Path 1 (which consists in the consecutive formation of all rungs by templating on previously misfolded structures) dominates over all others as soon as k_1_/k_2_ ≥ 4. Using Kramers’ theory and assuming comparable pre-factors, we find that the templating mechanism is the most prominent reaction pathway when the activation energies for single and double rung formations obey the relationship:

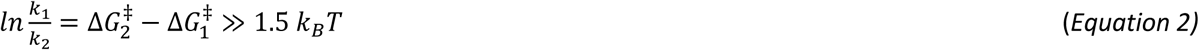

We expect this condition to be always satisfied. Indeed, processes 1 and 2 are characterized by the formation of the same number of hydrogen bonds, leading approximately to the same enthalpic gain. However, the process described by rate k_2_ requires the breaking of the double amount of native contacts, together with a two-fold entropy loss (compared to the single rung formation) when forming two de novo rungs. We can therefore conclude that the propagation reaction proceeds through a subsequent formation of individual rugs, using as template pre-existing 4RβS free end. We emphasize that our approach is likely to underestimate the dominance of the sequential misfolding mechanism. Indeed, it does not account for the direct cooperativity of hydrogen bonds and the long-range electrostatics favoring β-strand formation in presence of pre-formed β-sheets, as directly supported by previous computational and experimental evidence(26, 27).

The coarse-grained information about the reaction mechanism obtained by means of our theoretical model can be exploited to set up a fully atomistic rMD simulation of the PrP^C^ → PrP^Sc^ transition, using the 4RβS as a target structure (Fig. 4 and Supp. Movie). The 3D structure of moPrP^C^ (residues 105-230) was obtained by linking moPrP 121-231 (PDB 1XYX) to the adapted N-terminal fragment (residues 105-120) of huPrP^C^ (PDB 5yj5) which was mutated to the moPrP sequence. The initial state for the conversion simulation was generated by modifying a central dimer extracted from the tetrameric 4RβS at the end of 20 ns of restrained molecular dynamics simulation. The initial contact point between PrP^Sc^ and PrP^C^ was generated by leaving residues 89-104 of the C-terminal monomer anchored to the β-solenoid, which was then replaced by moPrP^C^ retaining the original disulfide bond. The rationale behind this modelling scheme derives from multiple previous reports indicating that the same region is a primary contact point between PrP^C^ and PrP^Sc^ (28), as well as by uncertainties regarding the state of the disulfide bond during PrP conformational transition. The complex was energy minimized in vacuum and then in explicit solvent in a dodecahedral box (with periodic boundary conditions and minimum wall distance of 20 Å) also containing six Cl^-^ ions to neutralize the charge of the system, using a steepest descent algorithm (protein topology was generated by Amber99SB-ILDN force field). The system was then equilibrated in the NVT ensemble (350 K) for 200 ps using the Nose-Hoover thermostat, and then in the NPT ensemble (350 K, 1 Bar) for 200 ps using the Nose-Hoover thermostat and the Parrinello-Rahman barostat. The model was then subjected to a modified protocol of rMD simulations, adapting the method to a sequential biasing. This scheme resulted in a progressive rMD simulation in which the structure of PrP^C^ was targeted to the growing solenoid in a rung-by-rung fashion. Target structures included an entire 4RβS solenoid and additional rungs at the growing C-terminus: Step 1, residues 89-115; Step 2, residues 89-151; Step 3, residues 89-190; Step 4, residues 89-230. In this rMD scheme, the equation of motion is modified by an history-dependent biasing force 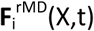, defined as:

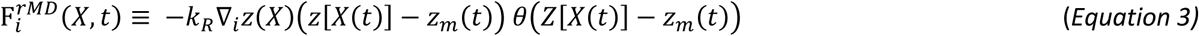

where *k*_R_ is the ratchet spring constant determining the strength of the biasing force applied on the system, in this case 10^-2^ kJ/mol. θ function is equal to 1 if its argument is positive, its value its 0 otherwise. *Z*[X(t)] is a collective coordinate defined as:

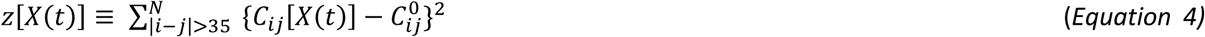

where *C*_ij_[X(t)] is the instantaneous contact map and *C*_ij_^0^ is the contact map of the target state. Their entries are chosen to smoothly interpolate between 0 and 1, depending on the relative distance of the atoms *i* and *j*. The *z*_m_ function indicates the smallest value assumed by the reaction coordinate *z*[X(t)] up to time t and. The contact map *C*_ij_(X) is defined as:

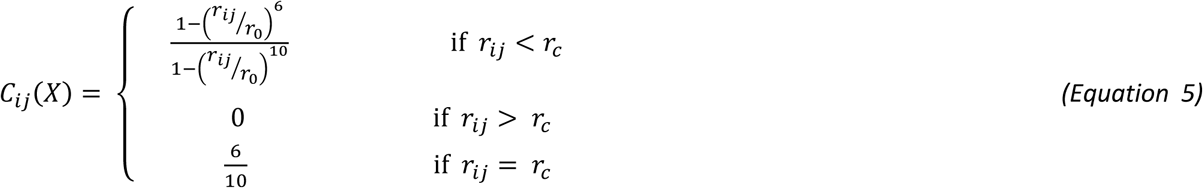

Where r_ij_ is the distance between the *i*-th and the *j*-th atom; r_0_ is a typical contact distance set to 7.5 Å; r_c_ is an upper limit to improve computational efficiency by excluding excessively distant atoms from the calculation (and it is set to 12.3 Å). In this way, no bias force acts on the system when the protein spontaneously proceeds towards the target state, while the external biasing force is only applied when the polypeptide tends to backtrack toward the initial state. We terminated each rMD simulation when the RMSD of the protein relative to the final state is stably lower than 0.8 Å. rMD simulations were performed in explicit solvent using Gromacs 4.6.5 with the plugin Plumed 2.0.2(29). Integration of motion was performed using a leap-frog algorithm with step equal to 2 fs. Temperature was maintained at 350 K (approximately PrP melting temperature) and pressure at 1 Bar using Nose-Hoover thermostat and the Parrinello-Rahman barostat. LINCS algorithm was applied to handle bonds constraints. Cut-off for short-range Coulomb and Van der Waals interactions was set to 10 Å, while long range electrostatics was treated using Particle Mesh Ewald.

### Data Analysis

Root Mean Squared Deviation of atomic positions (RMSD) was calculated “all-atoms” using Gromacs 5.1.2. The specific calculation of the β-solenoid/helix core RMSD involves all the following residues for the different systems:

4RβS Model: T94, H95, N96, Q97, K105, T106, N107, L108, K109, H110, V111, A112, G113, A114, A115, M128, L129, G130, S131, G141, N142, D143, W144, E145, D146, R147, Y148, Y149, R150, E151, D166, Q167, Y168, S169, N180, I181, T182, I183, K184, Q185, H186, T187, V188, T189, T190, T198, E199, T200, D201, T215, Q216, Y217, Q218, K219, E220, S221, Q222, A223, Y224, Y225.

Het-s Dimer: R225, N226, S227, A228, K229, D230, I231, R232, T233, E234, E235, R236, A237, R238, V239, Q240, L241, G242, V244, T261, N262, S263, V264, E265, T266, V267, V268, G269, K270, G271, E272, S273, R274, V275, L276, I277, G278, N279, E280.

LβH Model: G89, Q90, G91, G92, G93, T94, H95, N96, Q97, W98, N99, K100, N107, L108, K109, H110, V111, A112, G113, A114, A115, A116, A117, G118, A119, V120, V121, G122, G123, L124, G125, G126, T127, M128, L129, G130, S131, A132, M133, S134, R135, P136, M137, I138, H139, F140, G141, N142, D143, W144, E145, D146, D166, Q167, Y168, S169.

Secondary structure content and fraction of native/misfolded contacts were obtained using the Timeline and the Trajectory tools in VMD 1.9.2. Fraction of native and misfolded contacts are defined in the following equation

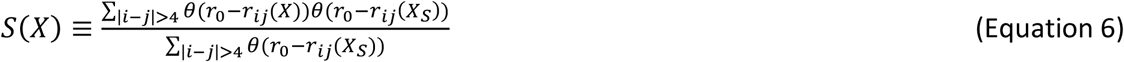

where r_ij_ is the distance between the *i*-th and *j*-th C_α_ atoms, r_0_ is the typical distance defining a contact between two residues and it is set at 7.5 Å, *X* is the instantaneous configuration of the protein during the simulation, *X_S_* is the atomic configuration of the protein in the reference state, that is the native PrP^C^ structure for the native contacts calculation, and the PrP^Sc^ structure for the misfolded contacts calculations. This quantity is evaluated only for the converting monomer.

Initial and final values of RMSD and secondary structure content were calculated by averaging the values over the last 5ns (of restrained or unrestrained MD) and then calculating the mean and standard deviation for the three trajectories. Inter-monomer distances in the tetramers are calculating using Chimera as an average of the distances between the mid-residue of each β-strand and the same residue in the first or the second next monomer. In particular, for β-strand-1: N96, G130, Y168, T200; for β-strand-2: N107, D143, T192, Y217; for β-strand-3: G113, Y149, V188, A223. Tetramer volumes were calculated using Chimera. Molecular grapics images were produced using the UCSF Chimera package from the Computer Graphics Laboratory, while graphs are plotted using Gnuplot.

## Supp. figure legends

**Supp. Fig. 1.**
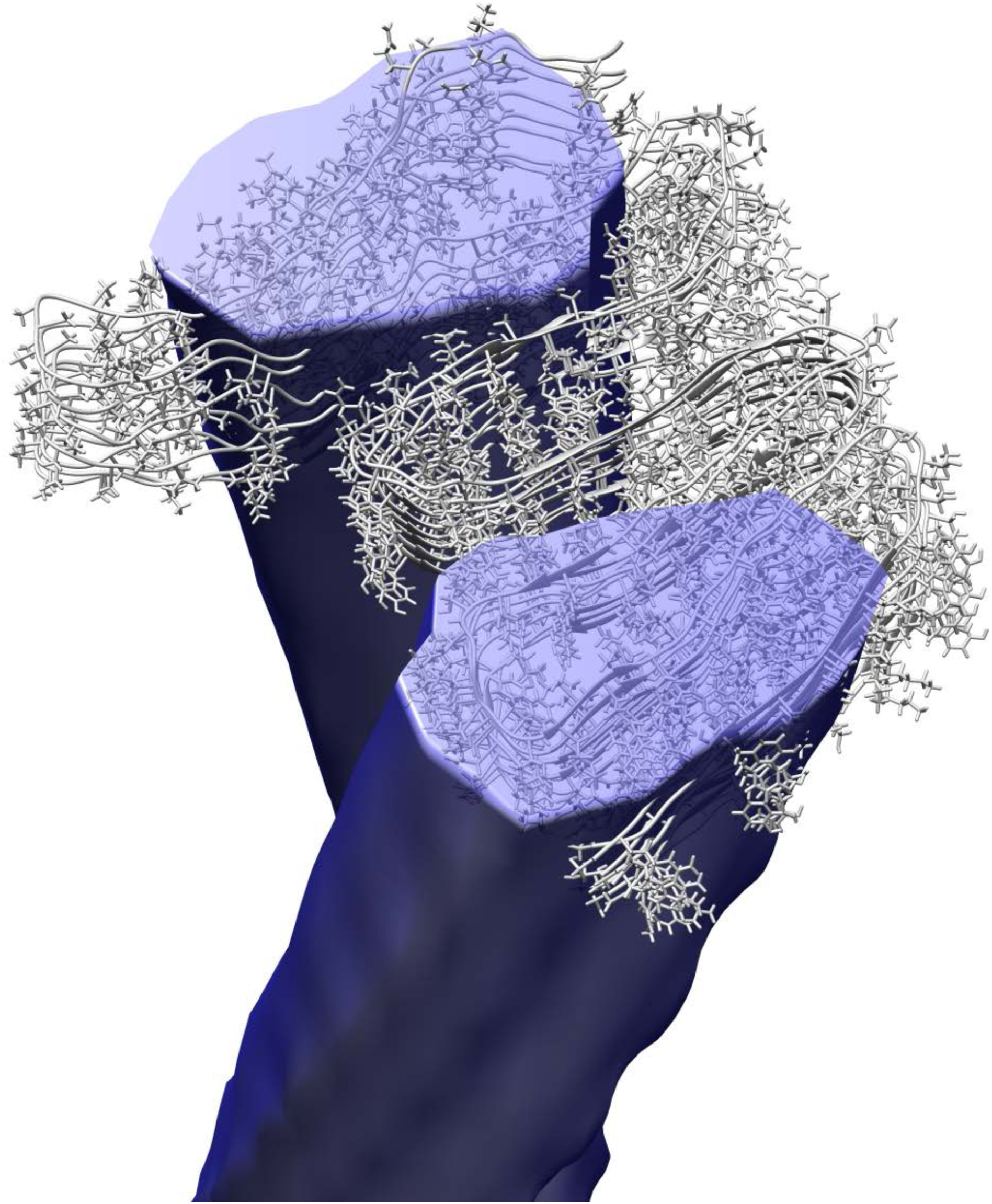
PIRIBS model of PrP^Sc^ superimposed to the cryo-EM map. The picture illustrates that the size of PIRIBS (80 Å x 50 Å) does not fit the fibril cross section (50 Å x 30 Å) obtained by cryo-EM (level of contouring equal to 3.05 in order to match the measured fibril diameter).

**Supp. Fig. 2.**
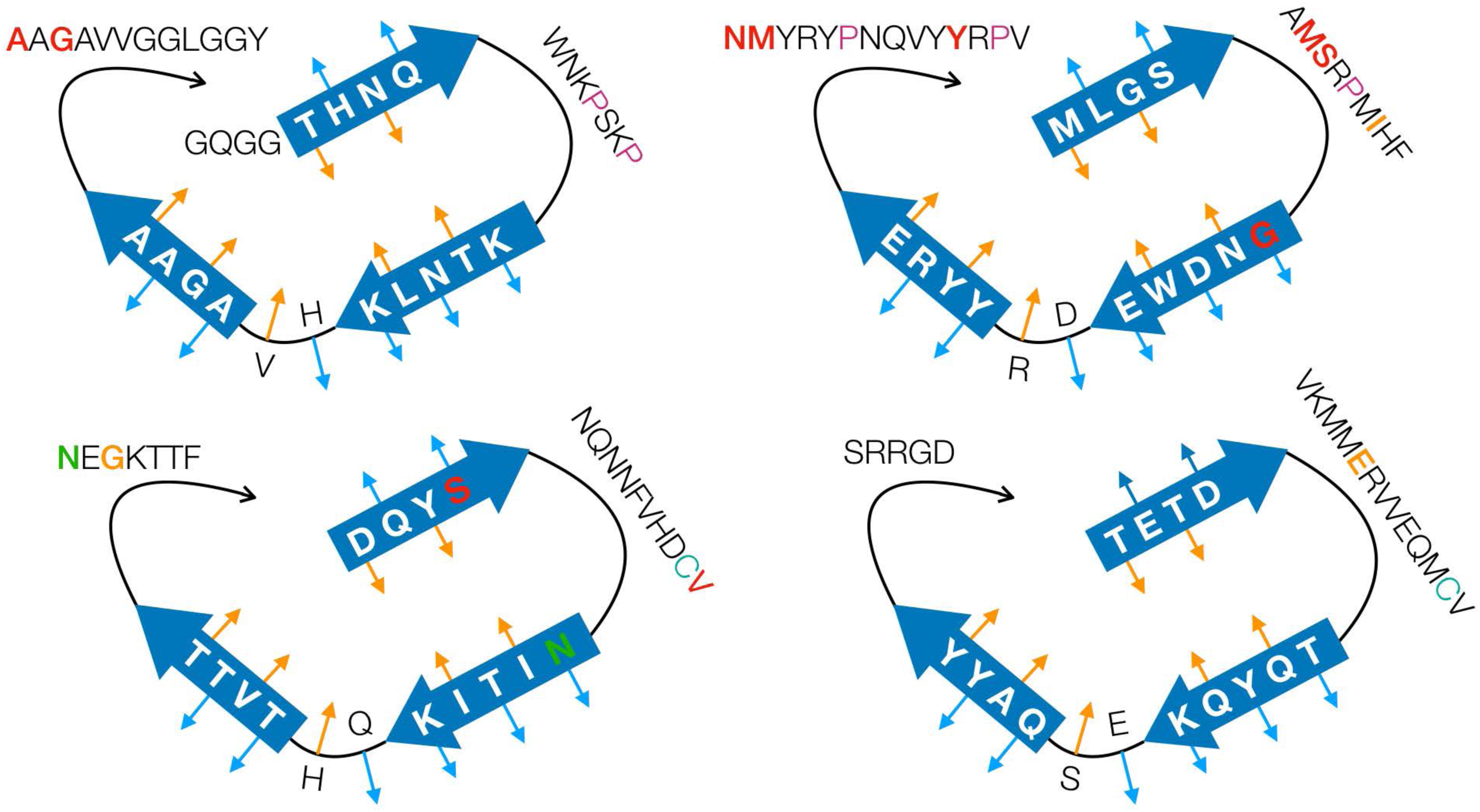
2D threading scheme of moPrP residues 89-230. 2D arrangement based on the general architecture of right-handed β-solenoid proteins with L-shaped cross section. Blue thick arrows indicate β-strands, thin cyan and thin orange arrows indicate the sidechain orientation (toward the solvent or toward the hydrophobic core, respectively). The scheme is used to thread the moPrP 89-230 sequence by considering different constraints. PK cleavage sites identified by mass spectrometry in two different previous reports are colored in red or orange. Glycosylation sites are labelled in green. Prolines are colored in purple. Cystine is indicated in cyan.

**Supp. Fig. 3.**
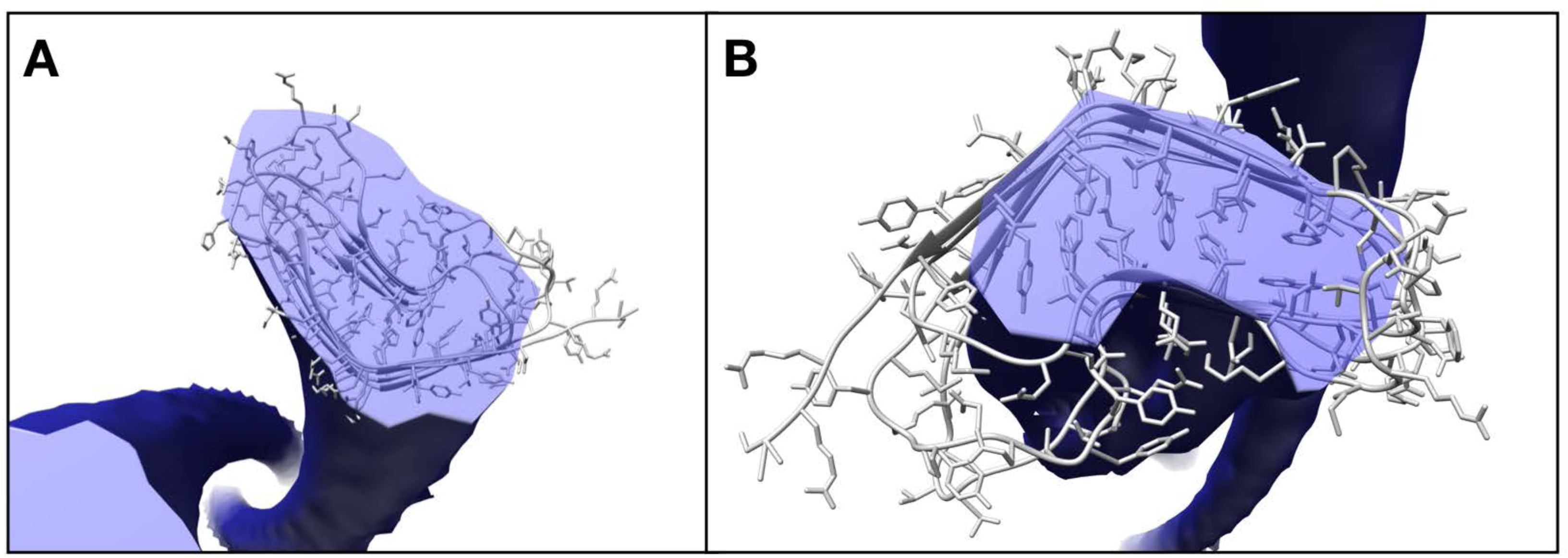
4RβS model of PrP^Sc^ superimposed to the cryo-EM map. The pictures illustrate that the size of 4RβS fits with the fibril cross section (50 Å x 30 Å) obtained by cryo-EM. **a**, Level of contouring was set to 3.05 to match the measured fibril diameter. **b**, Level of contouring was set to 3.60 to highlight the superimposition of the hydrophobic core of the 4RβS with the highest electronic density region of the fibril.

**Supp. Fig. 4.**
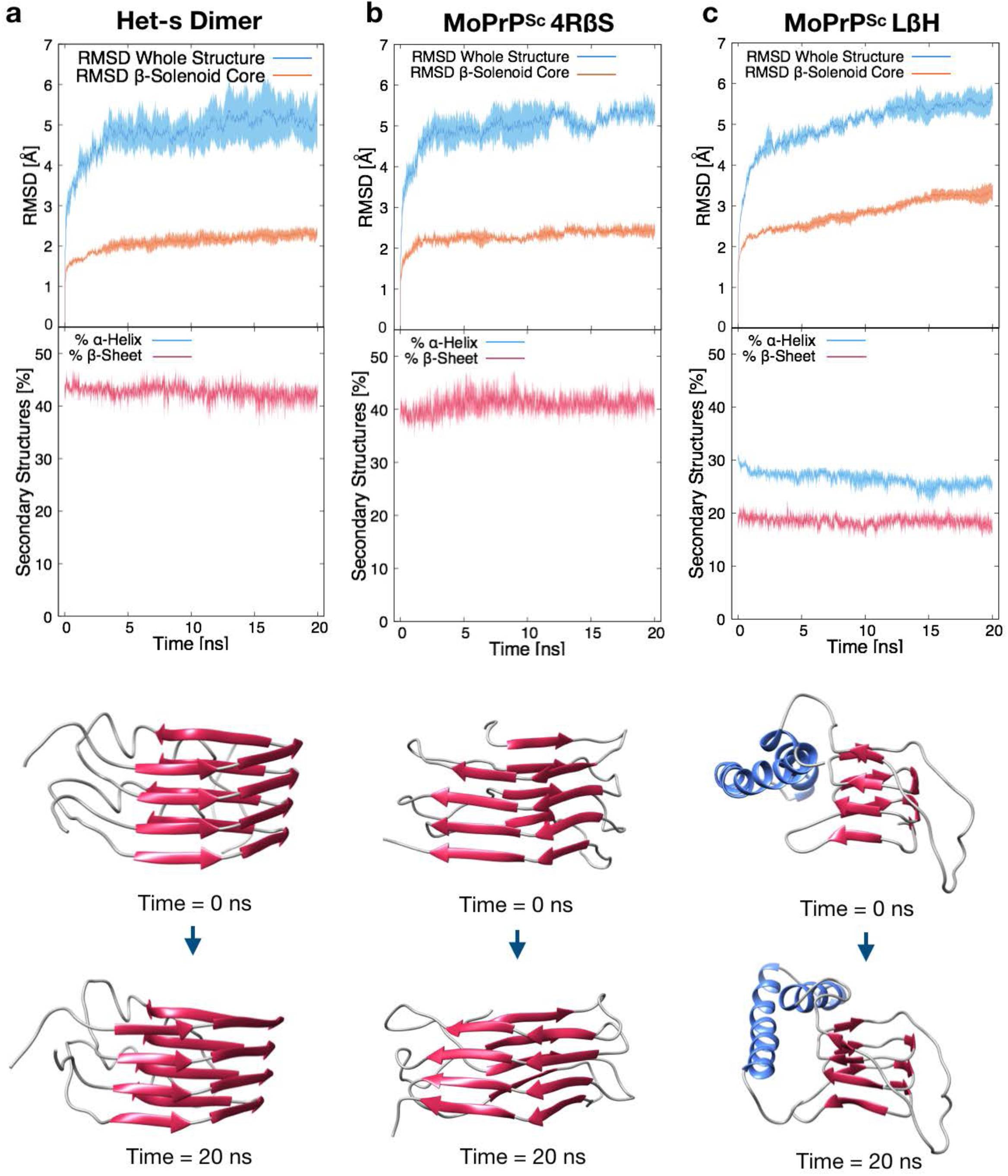
Summary of restrained MD simulations of the Het-s dimer, 4RβS and LβH PrP^Sc^. Upper graphs report the RMSD deviation from initial configurations for the entire structure (blue lines) or the β-solenoid core (orange lines) of the different proteins during the 20 ns of restrained MD simulations. Filled curves represents standard error of the mean. The graphs indicate a minor rearrangement of the β-solenoid core that is almost identical for the Het-s dimer and the 4RβS model, characterized by a RMSD of 2.3 ± 0.2 Å and 2.4 ± 0.2 Å respectively (calculated as the average of the three trajectories over the last 5 ns, ± standard deviation). In contrast, the hydrophobic core of the LβH model displays a higher RMSD deviation (3.3 ± 0.3 Å). Lower graphs indicate the α-helical (blue lines) or β-sheet (red lines) content of each protein. The latter is stable for all the three structures, likely due to the presence of the restraining potential for the whole length of the three simulations.

**Supp. Fig. 5.**
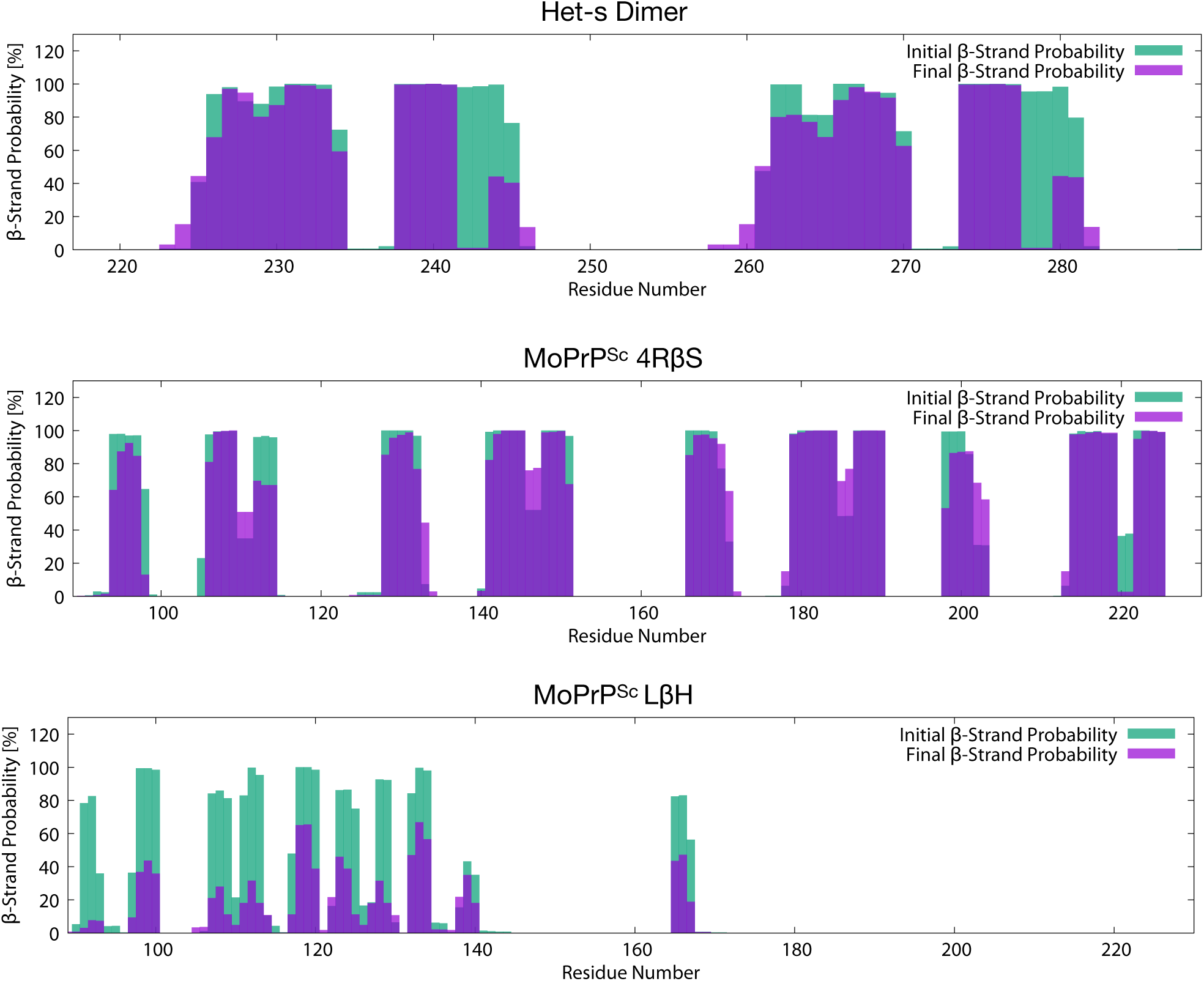
Probability distribution of β-sheet content for the Het-s dimer, 4RβS and LβH PrP^Sc^. Graphs show the probability of each residue to be in an extended conformation at the beginning of the simulation (green) or at the end of the simulation (purple) for the Het-s dimer, 4RβS and LβH PrP^Sc^ models. Initial or final probabilities were calculated from the last 5 ns of the restrained or unrestrained simulations, respectively.

**Supp. Fig. 6.**
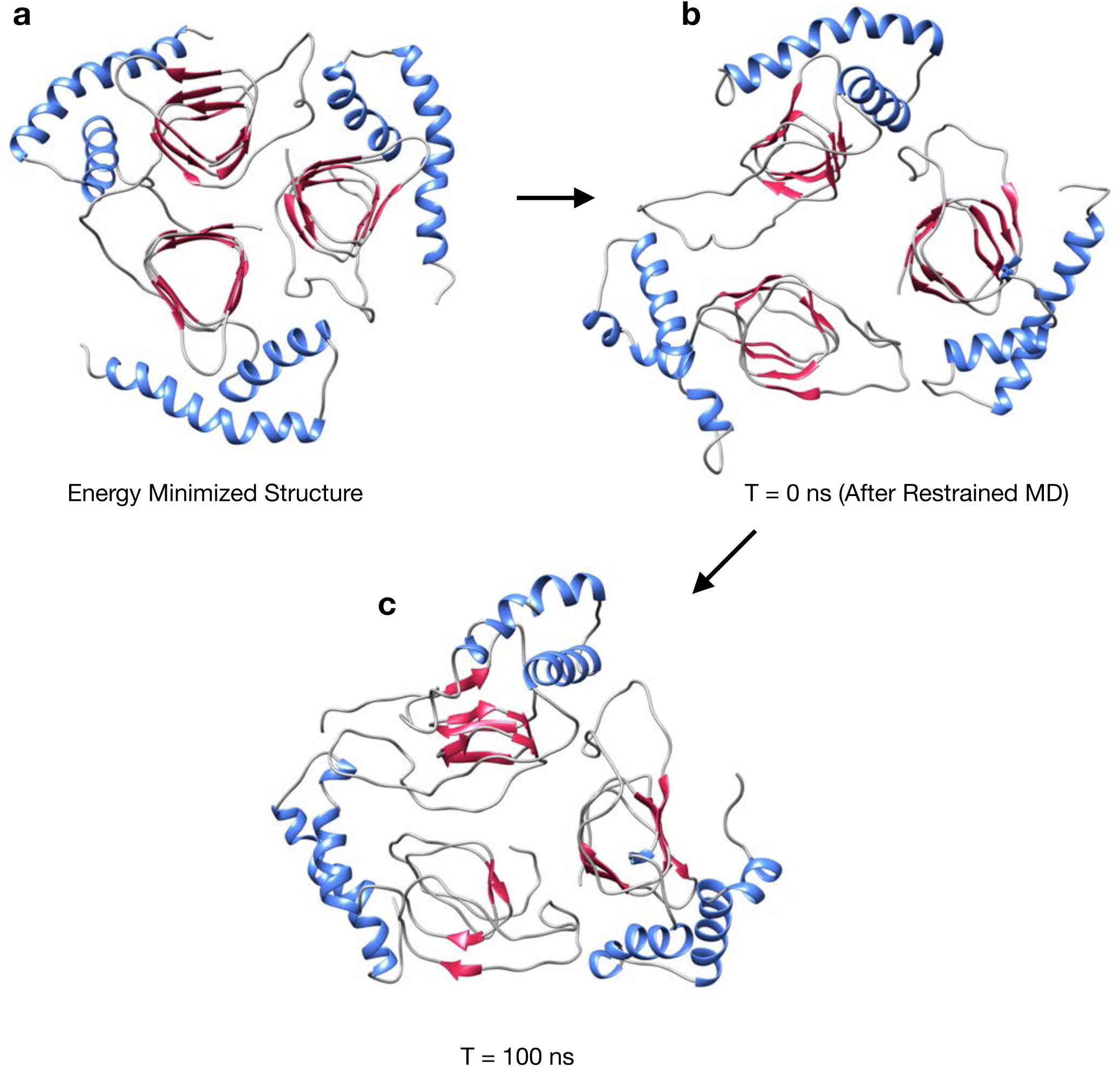
Initial and final conformations of LβH PrP^Sc^ after MD simulations. Pictures show the energy minimized structure (1), the same structure at end of the restrained (2) or unrestrained (3) MD simulations of the LβH model.

**Supp. Fig. 7.**
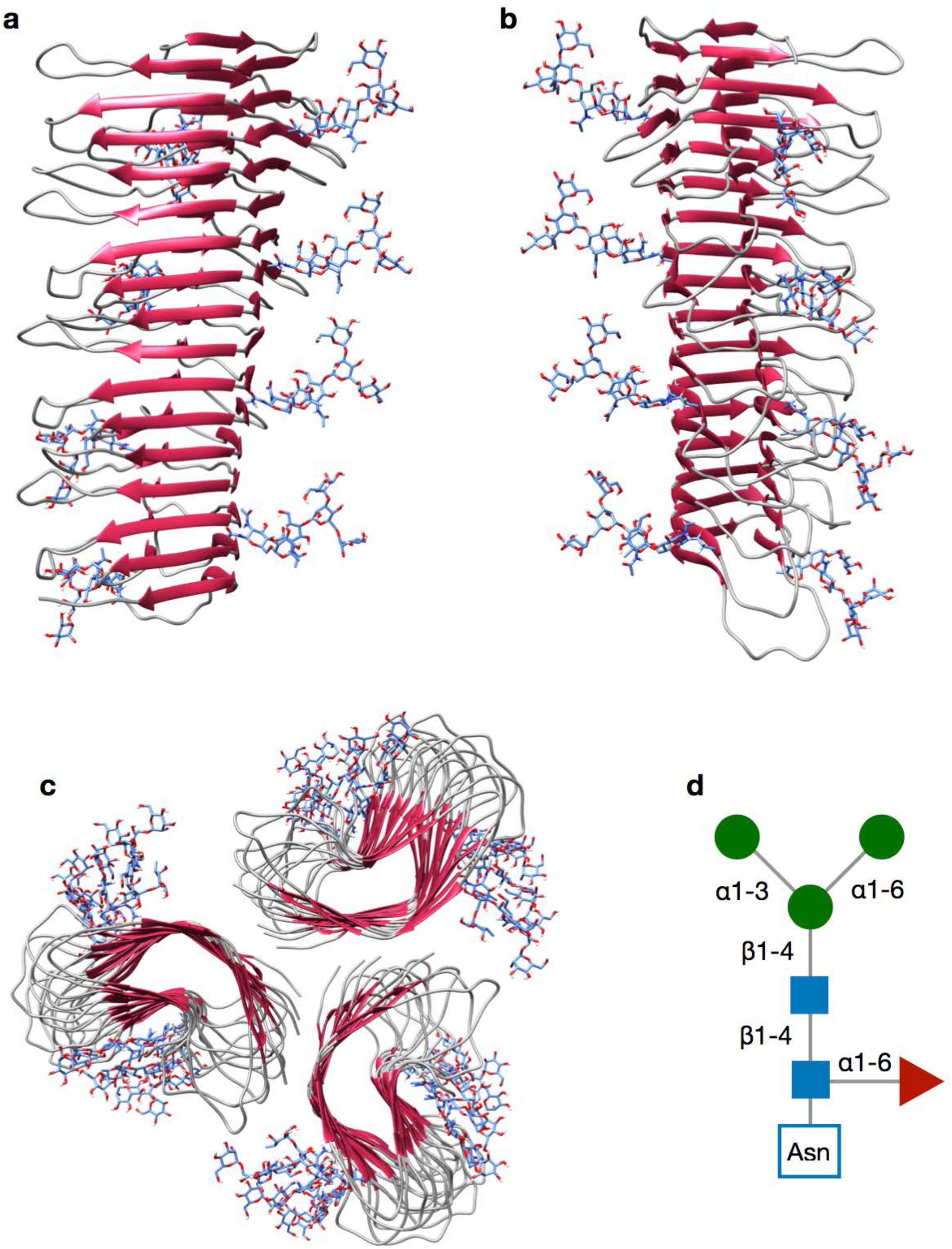
Illustration of the energy minimized 4RβS tetramer carrying complex glycans. A model of energy minimized tetrameric 4RβS structure carrying glycan residues is depicted in two different orientations (a, b). Top view of a laterally stacked trimer of glycosylated 4RβS (c), compatible with the 2D crystals diffraction data. A scheme of the complex glycan precursor added is depicted in (d), blue squares indicate N-acetylglucosamine, green circles indicate mannose and the red triangle indicate fucose.

**Supp. Fig. 8.**
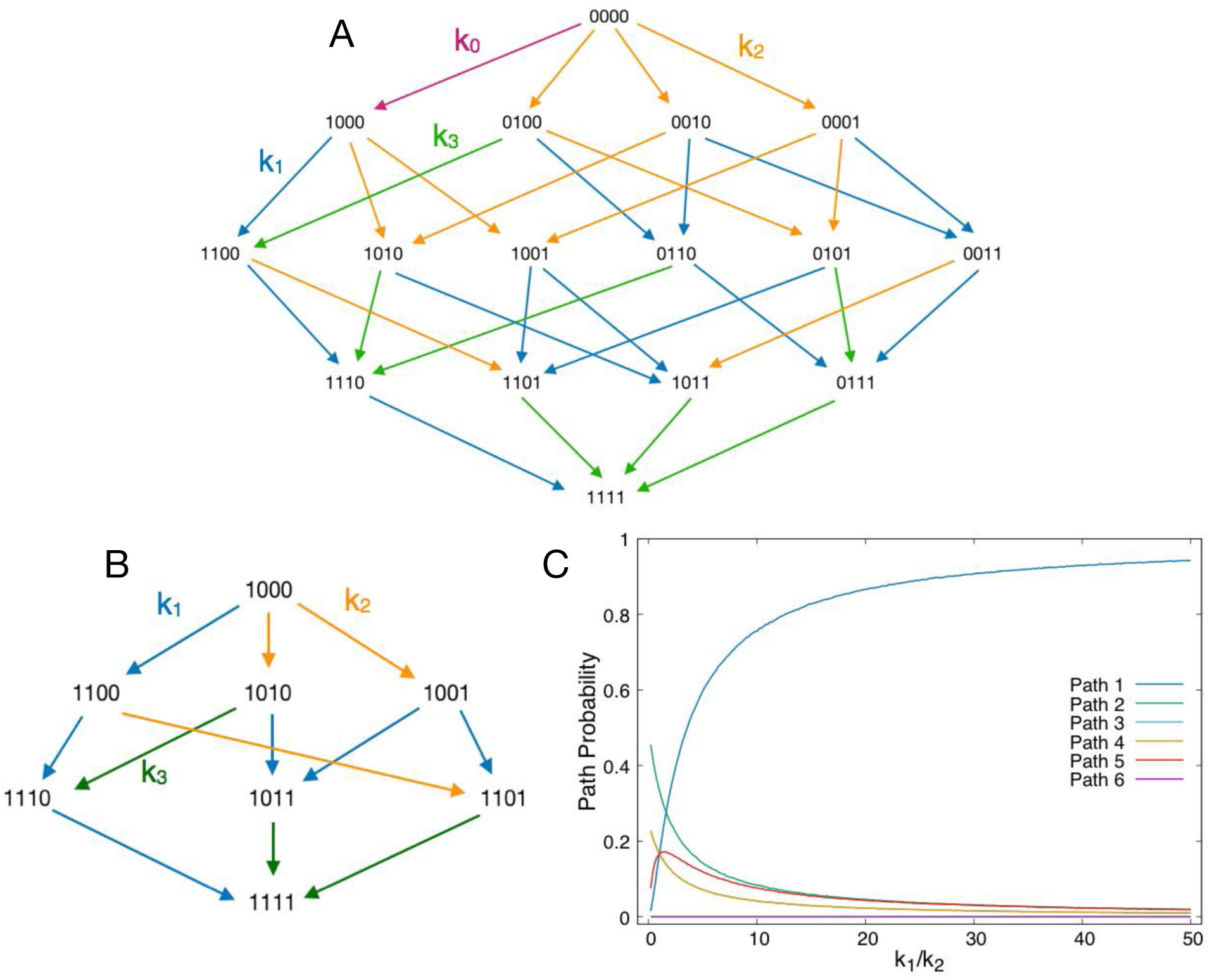
Statistical model for prion propagation mechanism. (a) Schematic representation of the network of transitions leading to the incorporation of PrP^C^ into PrP^Sc^. The purple arrow describes the formation of a single rung by templating the unstructured region of PrPC (89-115) on PrPSc (rate k0). The blue arrows describe the formation of a single by means of a templating process which involves the breaking the native contacts and the docking onto a pre-formed rung (rate k1). The orange arrows indicate the spontaneous formation and docking of two de-novo rungs (rate k2). Finally, the green arrows indicate the docking of preformed rungs belonging to two adjacent misfolded regions of the same chain (rate k3). (b) Simplified version of the network assuming as a priming reaction step the formation of the first rung. (c) Relative probability of the 6 reaction pathways as a function of the k_1_/k_2_ rate ratio.

**Supp. Movie. Full-atomistic model of prion propagation.** Visualization of the rMD simulation reconstructing the entire refolding events of PrP^C^ onto the 4RβS PrP^Sc^ model.

## Acknowledgements

HW acknowledges support through grant 201600029 from the Alberta Prion Research Institute. JRR was funded by grants BFU2017-86692-P and BFU2013-48436-C2-1-P from the Spanish Ministries of Economy and Competitiveness, and Education and Science, respectively, both partially including FEDER funds from the European Union. Calculations were performed using supercomputing facilities Finis Terrae II (CESGA, Spain) and Marconi (CINECA, Italy). This work was supported by a grant from Fondazione Telethon (Italy, TCP14009). GS is a recipient of a fellowship from Fondazione Telethon. EB is an Assistant Telethon Scientist at the Dulbecco Telethon Institute.

## Author contributions

JRR, PF, EB, HW and GS conceived and developed the project; JRR, AMS, GS and MR designed and built the 4RβS PrP^Sc^ model; GS and MR performed MD simulations and analyses; EB, PF, SO, GS and MR developed methods to reconstruct PrP misfolding; EB, GS, MR, PF and JRR wrote the original version of the manuscript, which was revised and accepted by all the authors.

## References

1. Prusiner SB (1982) Novel proteinaceous infectious particles cause scrapie. Science 216(4542):136–144.

2. Gremer L, et al. (2017) Fibril structure of amyloid-beta(1-42) by cryo-electron microscopy. Science 358(6359):116–119.

3. Wasmer C, et al. (2008) Amyloid fibrils of the HET-s(218-289) prion form a beta solenoid with a triangular hydrophobic core. Science 319(5869):1523–1526.

4. Vazquez-Fernandez E, et al. (2016) The Structural Architecture of an Infectious Mammalian Prion Using Electron Cryomicroscopy. PLoS Pathog 12(9):e1005835.

5. Wille H, et al. (2009) Natural and synthetic prion structure from X-ray fiber diffraction. Proc Natl Acad Sci U S A 106(40):16990–16995.

6. Prusiner SB (1998) The prion diseases. Brain Pathol 8(3):499–513.

7. Zahn R, et al. (2000) NMR solution structure of the human prion protein. Proc Natl Acad Sci U S A 97(1):145–150.

8. Requena JR & Wille H (2014) The structure of the infectious prion protein: experimental data and molecular models. Prion 8(1):60–66.

9. Govaerts C, Wille H, Prusiner SB, & Cohen FE (2004) Evidence for assembly of prions with left-handed beta-helices into trimers. Proc Natl Acad Sci U S A 101(22):8342–8347.

10. Groveman BR, et al. (2014) Parallel in-register intermolecular beta-sheet architectures for prion-seeded prion protein (PrP) amyloids. J Biol Chem 289(35):24129–24142.

11. Baskakov IV & Katorcha E (2016) Multifaceted Role of Sialylation in Prion Diseases. Front Neurosci 10:358.

12. Welker E, Raymond LD, Scheraga HA, & Caughey B (2002) Intramolecular versus intermolecular disulfide bonds in prion proteins. J Biol Chem 277(36):33477–33481.

13. Vazquez-Fernandez E, et al. (2012) Structural organization of mammalian prions as probed by limited proteolysis. PLoS One 7(11):e50111.

14. Sevillano AM, et al. (2018) Recombinant PrPSc shares structural features with brain-derived PrPSc: Insights from limited proteolysis. PLoS Pathog 14(1):e1006797.

15. Tiana G & Camilloni C (2012) Ratcheted molecular-dynamics simulations identify efficiently the transition state of protein folding. J Chem Phys 137(23):235101.

16. Lindorff-Larsen K, et al. (2010) Improved side-chain torsion potentials for the Amber ff99SB protein force field. Proteins 78(8):1950–1958.

17. S AB, Fant L, & Faccioli P (2015) Variational scheme to compute protein reaction pathways using atomistic force fields with explicit solvent. Phys Rev Lett 114(9):098103.

18. Ianeselli A, et al. (2018) Atomic Detail of Protein Folding Revealed by an Ab Initio Reappraisal of Circular Dichroism. J Am Chem Soc 140(10):3674–3682.

19. Bartolucci G, Orioli S, & Faccioli P (2018) Transition path theory from biased simulations. J Chem Phys 149(7):072336.

20. Nicholson EM, Mo H, Prusiner SB, Cohen FE, & Marqusee S (2002) Differences between the prion protein and its homolog Doppel: a partially structured state with implications for scrapie formation. J Mol Biol 316(3):807–815.

21. Kajava AV & Steven AC (2006) Beta-rolls, beta-helices, and other beta-solenoid proteins. Adv Protein Chem 73:55–96.

22. Pettersen EF, et al. (2004) UCSF Chimera--a visualization system for exploratory research and analysis. J Comput Chem 25(13):1605–1612.

23. Sali A & Blundell TL (1993) Comparative protein modelling by satisfaction of spatial restraints. J Mol Biol 234(3):779–815.

24. Berendsen HJC, Vanderspoel D, & Vandrunen R (1995) Gromacs - a Message-Passing Parallel Molecular-Dynamics Implementation. Comput Phys Commun 91(1-3):43–56.

25. Emsley P, Lohkamp B, Scott WG, & Cowtan K (2010) Features and development of Coot. Acta Crystallogr D 66:486–501.

26. Zhao YL & Wu YD (2002) A theoretical study of beta-sheet models: is the formation of hydrogen-bond networks cooperative? J Am Chem Soc 124(8):1570–1571.

27. Kung VM, Cornilescu G, & Gellman SH (2015) Impact of Strand Number on Parallel beta-Sheet Stability. Angew Chem Int Ed Engl 54(48):14336–14339.

28. Moroncini G, et al. (2004) Motif-grafted antibodies containing the replicative interface of cellular PrP are specific for PrPSc. Proc Natl Acad Sci U S A 101(28):10404–10409.

29. Bonomi M, et al. (2009) PLUMED: A portable plugin for free-energy calculations with molecular dynamics. Comput Phys Commun 180(10):1961–1972.

30. Friedman R & Caflisch A (2014) Wild type and mutants of the HET-s(218-289) prion show different flexibility at fibrillar ends: a simulation study. Proteins 82(3):399–404.

31. Mizuno N, Baxa U, & Steven AC (2011) Structural dependence of HET-s amyloid fibril infectivity assessed by cryoelectron microscopy. Proc Natl Acad Sci U S A 108(8):3252–3257.

